# Microbial glycoside hydrolases as antibiofilm agents with cross-kingdom activity

**DOI:** 10.1101/113696

**Authors:** Brendan D. Snarr, Perrin Baker, Natalie C. Bamford, Yukiko Sato, Hong Liu, Melanie Lehoux, Fabrice N. Gravelat, Hanna Ostapska, Shane R. Baistrocchi, Robert P. Cerone, Elan E. Filler, Matthew R. Parsek, Scott G. Filler, P. Lynne Howell, Donald C. Sheppard

**Affiliations:** Departments of Medicine, Microbiology and Immunology; Infectious Diseases in Global Health Program, Centre for Translational Biology, McGill University Health Centre. Montréal, Québec, H4A 3J1, Canada; Program in Molecular Structure & Function, Research Institute, The Hospital for Sick Children, Toronto, Ontario, M5G 1X8, Canada; Department of Biochemistry, University of Toronto, Toronto, Ontario, M5S 1A8, Canada; Los Angeles Biomedical Research Institute at Harbor-UCLA Medical Center, Torrance, CA, USA; Department of Microbiology, University of Washington, Seattle, Washington, USA; David Geffen School of Medicine at University of California, Los Angeles, CA, USA

**Keywords:** Biofilm, Aspergillus, Pseudomonas, Therapeutics, Exopolysaccharide

## Abstract

Galactosaminogalactan and Pel are cationic heteropolysaccharides produced by the opportunistic pathogens, *Aspergillus fumigatus* and *Pseudomonas aeruginosa*, respectively. These exopolysaccharides both contain 1,4-linked *N-*acetyl-D-galactosamine and play an important role in biofilm formation by these organisms. Proteins containing glycoside hydrolase domains have recently been identified within the biosynthetic pathway of each exopolysaccharide. Recombinant hydrolase domains from these proteins (Sph3_h_ from *A. fumigatus* and PelA_h_ from *P. aeruginosa*) were found to degrade their respective polysaccharides *in vitro*. We therefore hypothesized that these glycoside hydrolases could exhibit anti-biofilm activity, and further, given the chemical similarity between galactosaminogalactan and Pel, that they might display cross-species activity. Treatment of *A. fumigatus* with Sph3_h_ disrupted *A. fumigatus* biofilms with an EC_50_ of 0.4 nM. PelA_h_ treatment also disrupted pre-formed *A. fumigatus* biofilms with EC_50_ values similar to those obtained for Sph3_h_. In contrast, Sph3_h_ was unable to disrupt *P. aeruginosa* Pel-based biofilms, despite being able to bind to the exopolysaccharide. Treatment of *A. fumigatus* hyphae with either Sph3_h_ or PelA_h_ significantly enhanced the activity of the antifungals posaconazole, amphotericin B and caspofungin, likely through increasing antifungal penetration of hyphae. Both enzymes were non-cytotoxic and protected A549 pulmonary epithelial cells from *A. fumigatus-induced* cell damage for up to 24 hours. Intratracheal administration of Sph3_h_ was well tolerated, and reduced pulmonary fungal burden in a neutropenic mouse model of invasive aspergillosis. These findings suggest that glycoside hydrolases can exhibit activity against diverse microorganisms and may be useful as therapeutic agents by degrading biofilms and attenuating virulence.

**Significance:** The production of biofilms is an important strategy used by both bacteria and fungi to colonize surfaces and to enhance resistance to killing by immune cells and antimicrobial agents. We demonstrate that glycoside hydrolases derived from the opportunistic fungus *Aspergillus fumigatus* and Gram-negative bacterium *Pseudomonas aeruginosa* can be exploited to disrupt pre-formed fungal biofilms and reduce virulence. Additionally, these glycoside hydrolases can be utilized to potentiate antifungal drugs by increasing their hyphal penetration, to protect human cells from fungal-induced injury and to attenuate virulence of *A. fumigatus* in a mouse model of invasive aspergillosis. The findings of this study identify recombinant microbial glycoside hydrolases as promising therapeutics with the potential for anti-biofilm activity against pathogens across different taxonomic kingdoms.

## body

The mould *Aspergillus fumigatus* and the Gram-negative bacterium *Pseudomonas aeruginosa* are opportunistic pathogens that cause pulmonary infection in immunocompromised patients and individuals who suffer from chronic lung diseases such as cystic fibrosis and bronchiectasis. *A. fumigatus* is the second most common nosocomial fungal infection (1) and ~10% of all nosocomial bacterial infections are caused by *P. aeruginosa* (2). Mortality associated with *P. aeruginosa* infections is high (3), and has increased with the emergence of multi- and even pan-resistance to antibiotics (3, 4). Similarly, invasive aspergillosis is associated with mortality rates of up to 50% (5), and increasing rates of antifungal resistance have been reported worldwide (6). These factors underscore the urgent need for new effective therapies for these infections.

Although *A. fumigatus* and *P. aeruginosa* are members of different taxonomic kingdoms, both produce biofilms that constitute a protective lifestyle for the organism. Biofilms are complex communities of microorganisms that grow embedded in an extracellular matrix composed of DNA, protein, and exopolysaccharide (7). Biofilm formation provides a significant advantage to these organisms as the matrix mediates adherence to host cells (8, 9) and aids in the resistance to both antimicrobial agents (10, 11) and host immune defences (12, 13). *A. fumigatus* biofilm formation is dependent on the cationic polysaccharide galactosaminogalactan (GAG), a heteroglycan composed of α1,4-linked galactose and *N*-acetyl-D-galactosamine (GalNAc) that is partially deacetylated (14, 15). In comparison, *P. aeruginosa* has the genetic capacity to produce three biofilm exopolysaccharides; alginate, Psl and Pel (16). GAG shares several similarities with Pel, which has been identified as a cationic heteroglycan composed of 1,4-linked GalNAc and *N*-acetyl-D-glucosamine (GlcNAc) (17). Like GAG, the cationic nature of Pel results from partial deacetylation of the polymer (17). Most clinical and environmental isolates of *P. aeruginosa* utilize Pel and Psl during biofilm formation (18). Alginate is dispensable for biofilm formation and is only observed in chronic pulmonary infection when strains switch to a mucoid phenotype (18, 19).

Strains of *Aspergillus* and *P. aeruginosa* with impaired GAG, or Pel and Psl biosynthesis exhibit attenuated virulence (20, 21), suggesting that targeting these exopolysaccharides may be a useful therapeutic strategy. We previously demonstrated that recombinant glycoside hydrolases PelA_h_ and PslG_h_, encoded in the *pel* and *psl* operons of *P. aeruginosa*, respectively, target and selectively hydrolyze the Pel and Psl exopolysaccharide components of the *Pseudomonas* biofilm matrix (22). Treatment with these enzymes rapidly disrupts established biofilms, increasing the susceptibility of *P. aeruginosa* to human neutrophil killing and potentiation of the antibiotic colistin (22).

Our recent work on *Aspergillus* has identified a cluster of five genes, which encode the proteins necessary for GAG biosynthesis (15). As with *P. aeruginosa,* we found that the product of one of these genes contains a glycoside hydrolase domain, Sph3_h_, that is capable of hydrolyzing purified and cell wall-associated GAG (23). In the present study we assessed the therapeutic potential of Sph3_h_ in disrupting fungal biofilms. We establish that the exogenous addition of Sph3_h_ is capable of rapidly disrupting existing biofilms of this organism at nanomolar concentrations. Additionally, we demonstrate cross-kingdom activity, as the *P. aeruginosa* glycoside hydrolase, PelA_h_, was able to disrupt *A. fumigatus* biofilms. While Sph3_h_ was able to bind Pel, it was unable to disrupt pre-formed *P. aeruginosa* Pel-mediated biofilms. Treatment with Sph3_h_ or PelAh increased the susceptibility of wild-type and azole-resistant *A. fumigatus* strains to lipophilic antifungal drugs. Kinetic studies with labelled posaconazole indicate that the increased susceptibility to antifungals is due to increased penetration of fungal cells by these agents. Both Sph3_h_ and PelA_h_ were non-toxic to mammalian cells and protected epithelial cells from *A. fumigatus-induced* damage for up to 24 hours. Intratracheal delivery of Sph3_h_ was well tolerated by mice and significantly reduced the fungal burden of immunocompromised mice infected with *A. fumigatus*. Our results suggest that glycoside hydrolases have the potential to be effective anti-biofilm therapeutics that can mediate activity against evolutionarily diverse microorganisms.

## Results

### Sph3_h_ disrupts preformed *A. fumigatus* biofilms

Our previous work demonstrated that Sph3_h_ from *A. fumigatus* and *Aspergillus clavatus* can hydrolyze both purified and cell wall-bound GAG on young hyphae (23). We therefore sought to determine if the degradation of GAG by Sph3_h_ could disrupt established *A. fumigatus* biofilms. Treatment with Sph3_h_ for one hour disrupted established *A. fumigatus* biofilms with an effective concentration for 50% activity (EC_50_) of 0.45 ± 1.31 nM **(Fig. 1a)**. Biofilm disruption was associated with a marked reduction in hyphae-associated GAG as detected by lectin staining **(Fig. 1b and c)** and scanning electron microscopy **(Fig. 1d)**. A catalytic variant Sph3_h_ D166A, which does not mediate GAG hydrolysis (23), displayed a greater than 500-fold reduction in anti-biofilm activity **(Fig. 1a)** and failed to mediate degradation of biofilm-associated GAG **(Fig. 1b and c)**. Collectively, these data suggest that biofilm disruption is mediated through the enzymatic hydrolysis of GAG.

**Figure 1.**
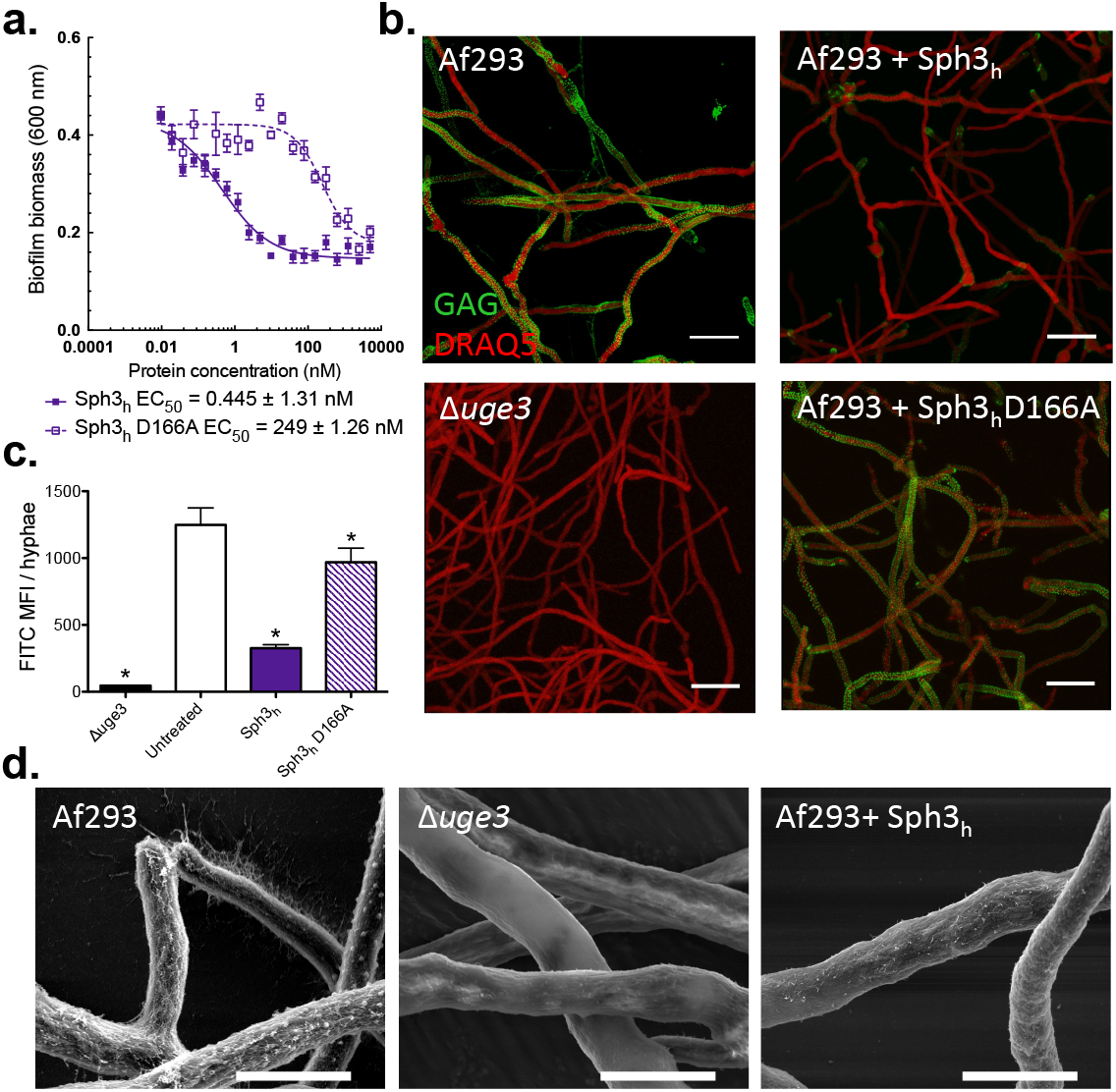
Treatment with Sph3_h_ disrupts *A. fumigatus.* biofilms and degrades GAG on the surface of hyphae. (a) Crystal violet staining of established *A. fumigatus* biofilms treated with the indicated concentration of each hydrolase. Each data point represents the mean of *n* = 20 with error bars indicating standard error (SE). EC_50_ indicates the 50% effective concentration ± SE. **(b)** Effects of the indicated hydrolases on cell wall associated GAG. Hyphae of the indicated strains grown in the absence of hydrolase treatment (left column) or following exposure to 0.5 μM of the indicated hydrolases (right column). Cell wall-associated GAG was visualized using FITC-conjugated lectin staining (green) with DRAQ5 as a counterstain (red). Scale bars = 20 μm. The GAG-deficient *Δuge3* mutant was included as a negative control. **(c)** Quantification of lectinstaining from panel (b). Each data point represents the mean fluorescence intensity of at least 7 hyphae with error bars indicating SE. * indicates a significant difference *(p* < 0.05) relative to the untreated *A. fumigatus* as determined by one-way ANOVA with Dunnett’s multiple comparison test. **(d)** Scanning electron micrographs of hyphae of the indicated strains grown in the absence of hydrolases (left & middle) and following exposure to 0.5 μM of Sph3_h_ (right column). Scale bars = 5 μm.

To validate that fungal biofilm disruption by Sph3_h_ is not restricted to the *A. fumigatus* laboratory strain Af293, the activity of Sph3_h_ was evaluated against four clinical *A. fumigatus* isolates. Sph3_h_ disrupted biofilms of all isolates tested at EC_50_ values < 0.15 nM **(Fig. S1)**. These results confirm the role of GAG in biofilm formation and indicate that Sph3_h_ exhibits anti-biofilm activity across a wide range of *A. fumigatus* strains.

### The bacterial hydrolase PelA_h_ hydrolyses GAG and disrupts fungal biofilms

Given that GAG and Pel are both cationic exopolysaccharides containing 1,4-linked GalNAc (14, 17), we hypothesized that PelA_h_ might exhibit activity against GAG. Consistent with this hypothesis, an *in vitro* reducing sugar assay demonstrated that PelA_h_ was capable of hydrolyzing purified GAG **(Fig. S2)**. Furthermore, using the crystal violet biofilm assay, we found that PelA_h_ disrupted *A. fumigatus* fungal biofilms with an EC_50_ value of 2.80 ± 1.14 nM **(Fig. 2a)**. The treatment of *A. fumigatus* hyphae with PelA_h_ also resulted in a reduction in the amount of cell wall-associated GAG **(Fig. 2b-d)** as was observed with Sph3_h_ treatment. The PelA_h_ catalytic variant, PelA_h_ E218A, which is markedly impaired in Pel hydrolysis and is inactive against *P. aeruginosa* biofilms (22) did not significantly hydrolyze GAG at concentrations as high as 12 μM **(Fig. S2)**. Consistent with this observation, PelA_h_ E218A was also several hundred-fold less active against *A. fumigatus* biofilms and did not degrade hyphae-associated GAG **(Fig. 2b and c)**. These results suggest that PelA_h_ disrupts *A. fumigatus* biofilms through the hydrolysis of biofilm associated GAG.

**Figure 2.**
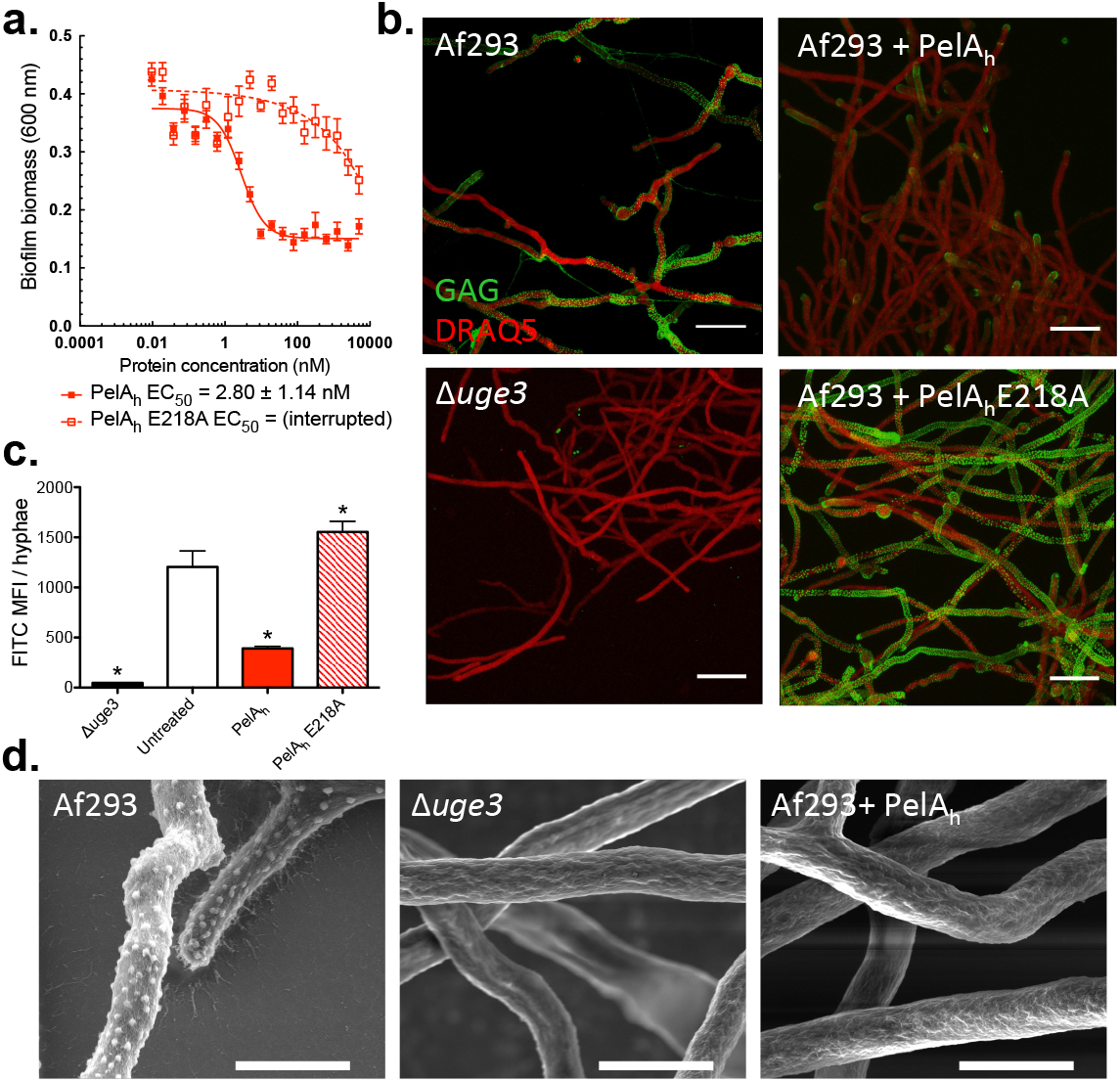
PelA_h_ disrupts *A. fumigatus* biofilms and degrades GAG. (a) Effects of PelA_h_ on *A. fumigatus* biofilms. Crystal violet staining of established *A. fumigatus* biofilms treated with the indicated concentration of PelA_h_ or PelA_h_ E218A. Each data point represents the mean of *n* = 20 with error bars indicating SE. EC_50_ reported ± SE. **(b)** Effects of the indicated hydrolases on cell wall-associated GAG. Established hyphae of the indicated strains were untreated (left column) or exposed to 0.5 μM of the indicated hydrolases (right column). Cell wall-associated GAG was detected using FITC-conjugated lectin staining (green), with DRAQ5 as a counterstain (red). Scale bars = 20 μ m. **(c)** Mean fluorescent intensity of lectin staining in panel (b). Each data point represents the mean of at least 7 hyphae with error bars indicating SE. The * indicates a significant difference *(p* < 0.05) relative to the untreated *A. fumigatus* as determined by one-way ANOVA with Dunnett’s multiple comparison test. **(d)** Scanning electron micrographs of hyphae of the indicated strains grown in the absence of hydrolase treatment (left & middle) or following treatment with 0.5 μM of PelA_h_ (right). Scale bars = 5 μm.

### Sph3_h_ binds Pel but does not disrupt established *P. aeruginosa* biofilms

Given that PelA_h_ can hydrolyze GAG and disrupt GAG-mediated biofilms, we hypothesized that Sph3_h_ may exhibit activity against Pel and Pel-mediated biofilms. The inability to purify sufficient quantities of Pel precluded us from utilizing it as a substrate. Therefore, to examine whether Sph3_h_ was capable of hydrolyzing Pel, the enzyme was exogenously applied to biofilms produced by the Pel overproducing *P. aeruginosa* strain PAO1 *ΔwspF Δpsl* P_BAD_*pel*. Treatment of these established biofilms with Sph3_h_ did not affect levels of Pel within the biofilms as visualized by lectin staining **(Fig. 3a and b)**, nor did it reduce biofilm biomass, even at concentrations exceeding 10 μM **(Fig. 3c)**.

**Figure 3.**
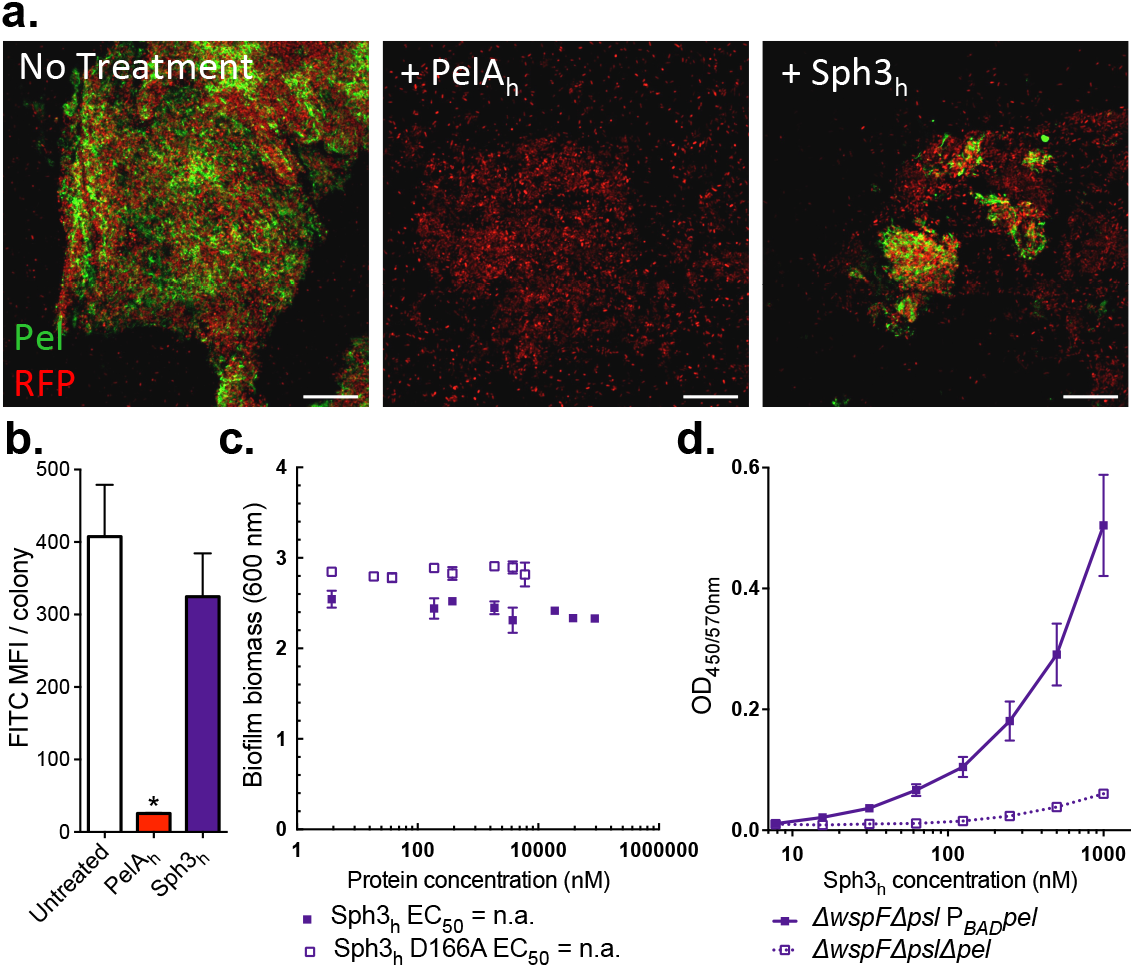
Sph3_h_ binds Pel but is inactive against established P *aeruginosa* biofilm. **(a)** Established biofilms of RFP-producing *P. aeruginosa* overexpressing the Pel operon (red) were untreated (left) or exposed to 0.5 μM of the indicated hydrolases (middle, right). Biofilm-associated Pel was detected using FITC-conjugated *Wisteria fluoribunda* lectin staining (green). Scale bars = 20 μm. **(b)** Mean fluorescent intensity of lectin staining in panel (a). Each data point represents the mean of at least 4 *P. aeruginosa* colonies with error bars indicating SE. * indicates a significant difference *(p* < 0.001) relative to the untreated *P. aeruginosa* as determined by 1-way ANOVA with Dunnett’s multiple comparison test. **(c)** Effects of Sph3_h_ on Pel-mediated *P aeruginosa* biofilms. Crystal violet staining of established biofilms of *P. aeruginosa* overexpressing the Pel operon incubated with the indicated concentrations of Sph3_h_ or Sph3_h_ D166A. Each data point represents the mean of *n* = 3 with error bars indicating SE. EC_50_ reported ± SE. **(d)** Sph3_h_ binding to Pel polysaccharide. Microtiter plates were coated with culture supernatants of the indicated *P aeruginosa* strains and the binding of Sph3_h_ was determined using an anti-Sph3_h_ antibody. Each data point represents the mean of 3 independent experiments with error bars indicating SE.

Since Sph3_h_ did not hydrolyze Pel within *P. aeruginosa* biofilms, we tested whether the enzyme was capable of recognizing and binding this polysaccharide. Using an ELISA-based binding assay we observed dose-dependent binding of Sph3_h_ to culture supernatants from the Pel over-producing *P. aeruginosa* strain, but not from supernatants of the Pel-deficient strain PAO1 *ΔwspF Δpel Δpsl* **(Fig. 3d)**. These data suggest that the inability of Sph3_h_ to disrupt Pel-mediated biofilms is likely a consequence of an inability to hydrolyze Pel rather than being unable to bind the polysaccharide. Dose-dependent binding of the inactive Sph3_h_ D166A variant to GAG-containing culture supernatants was also observed **(Fig. S3)**, suggesting that binding of hydrolases to exopolysaccharides is insufficient to disrupt established biofilms in the absence of enzymatic cleavage of the polymer.

### Sph3_h_ and PelA_h_ potentiate antifungals by enhancing their intracellular penetration

The Pel polysaccharide enhances resistance to several antibiotics including aminoglycosides and colistin (22, 24,25). Since biofilm formation by *A. fumigatus* is associated with increased resistance to a number of antifungal agents (26–28), we hypothesized that GAG may have an analogous function to Pel in enhancing resistance to antifungal agents. To test this hypothesis, we investigated whether Sph3_h_ or PelA_h_ could potentiate the activity of commonly used antifungal drugs. Treatment of established fungal biofilms with either enzyme resulted in a significant reduction in the MIC_50_ of the azole posaconazole, the polyene amphotericin B, and the echinocandin caspofungin **(Fig. 4a)**. Sph3_h_ or PelA_h_ treatment produced a similar increase in sensitivity to posaconazole for both azole-sensitive and azole-resistant strains of *A. fumigatus* **(Fig. S4)**. Susceptibility to voriconazole, a smaller and more polar azole, was unaffected by treatment with either glycoside hydrolase **(Fig. 4a)**. Since both posaconazole and voriconazole have the same intracellular target, these findings suggest that cationic GAG mediates antifungal resistance by hindering cellular uptake of large, nonpolar molecules such as posaconazole. To investigate this hypothesis, the effect of Sph3_h_ on intracellular penetration of posaconazole was examined using posaconazole conjugated to the fluorophore BODIPY (BDP-PCZ). Previous work has established that BDP-PCZ displays similar cellular and subcellular pharmacokinetics to unmodified posaconazole (29). Fluorometric studies revealed that Sph3_h_-treatment resulted in higher accumulation of BDP-PCZ within *A. fumigatus* hyphae **(Fig. 4b)**. This finding indicates that GAG protects *A. fumigatus* from the action of lipophilic antifungals by limiting their penetration into hyphae.

**Figure 4.**
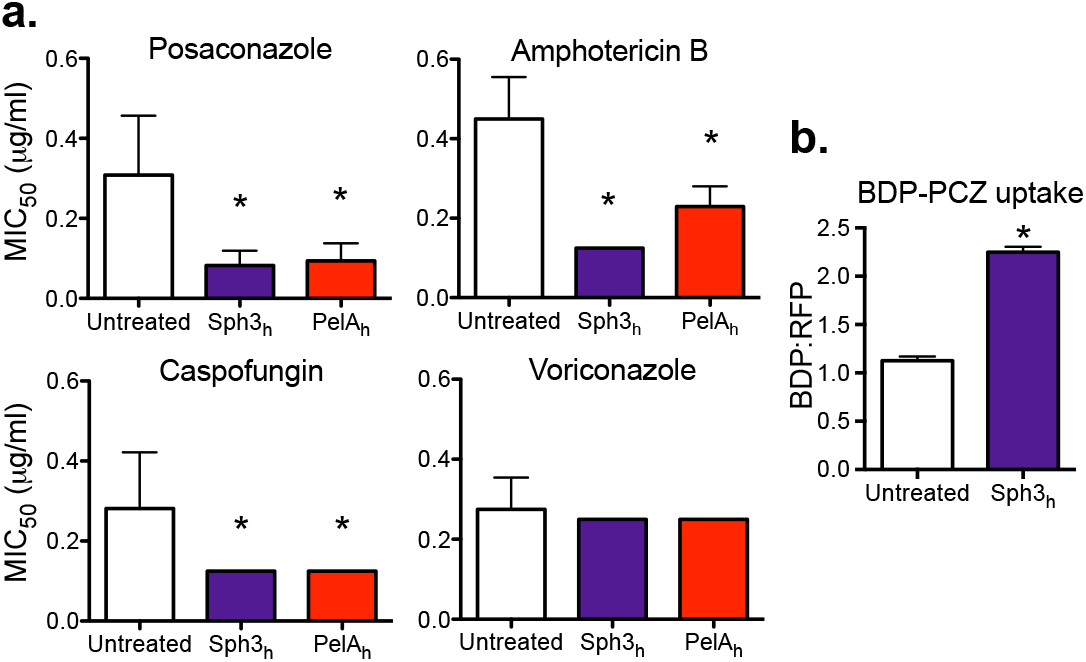
Glycoside hydrolases increase sensitivity of *A. fumigatus* to antifungal agents. **(a)** Established biofilms of wild-type *A. fumigatus* strain Af293 were treated with the indicated concentrations of antifungals with or without 0.5 μM of the indicated hydrolase and the viability of the resulting biofilms was then measured using the XTT metabolic assay. Susceptibility to antifungals was quantified by determining the antifungal concentration resulting in a 50% reduction in fungal metabolic activity (MIC_50_) as compared to untreated controls. Bars represent the mean of at least *n* = 4 with error bars indicating SE. **(b)** Effects of hydrolase therapy on antifungal uptake. *A. fumigatus* hyphae were treated with 1 μM Sph3_h_ then exposed to 2 μg/ml BDP-PCZ. Uptake of BDP-PCZ was quantified via fluorometry. Each bar represents the mean of 3 independent experiments with error bars indicating SE. The * indicates a significant difference *(p* < 0.05) relative to untreated control samples as determined by 1-way ANOVA with Dunnett’s multiple comparison test in (a), or two-tailed student’s T-test in (b).

### Recombinant Sph3_h_ and PelA_h_ protect epithelial cells from damage by *A. fumigatus*

*A. fumigatus* GAG-mediated adherence is required for *A. fumigatus* to damage A549 pulmonary epithelial cells *in vitro* (20). We therefore tested whether treatment with either Sph3_h_ or PelA_h_ could protect epithelial cells from fungal-induced injury using an established chromium (^51^Cr) release damage assay (30). We first established that the enzymes were not cytotoxic and that the addition of Sph3_h_ or PelA_h_ to uninfected A549 cell monolayers did not cause detectable cellular damage **(Fig. S5a)**, a finding verified with the IMR-90 human lung fibroblast cell line **(Fig. S5b)**. These data are consistent with the lack of cytotoxicity previously reported for PelA_h_ (22). Next, we assessed whether Sph3_h_ or PelA_h_ were able to protect A549 cell monolayers from damage by *A. fumigatus.* Sph3h reduced epithelial cell injury by > 80% for 24 hours **(Fig. 5a)**. Treatment with PelA_h_ also protected epithelial cells from *A. fumigatus-induced* damage **(Fig. 5a)**. The protective effect of PelA_h_ was shorter than that observed with Sph3_h_, and was lost before 24 hours of treatment. The addition of protease inhibitors extended PelA_h_- mediated epithelial cell protection to 24 hours **(Fig. S5c)**, suggesting that the decrease in PelA_h_ mediated protection was likely due to proteolytic degradation of the recombinant protein. Epithelial cell protection was not observed with the catalytic variants, PelA_h_ E218A or Sph3_h_ D166A, suggesting that the hydrolytic activity of the enzymes is required for protection **(Fig 5a)**.

**Figure 5.**
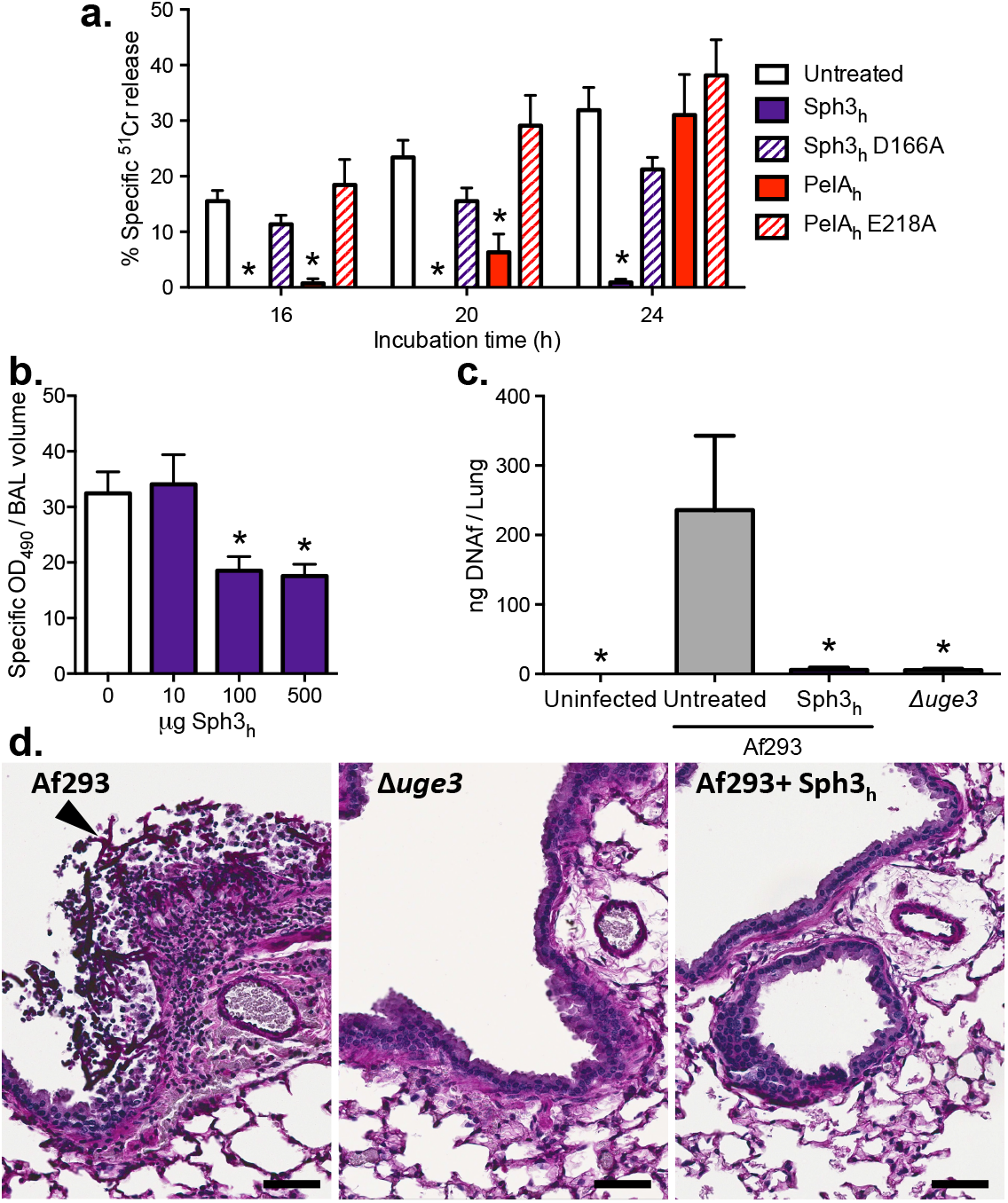
Effects of hydrolases on *A. fumigatus-induced* airway epithelial cell damage and *in vivo* pulmonary infection. **(a)** ^51^Cr-loaded A549 pulmonary epithelial cells were incubated with conidia of wild-type *A. fumigatus* in the presence or absence of 0.5 μM concentrations of the indicated hydrolases. Epithelial cell damage was determined by measurement of the amount of ^51^Cr released into supernatant at the indicated time points. Each bar represents the mean of at least 3 (Untreated, Sph3_h_, Sph3_h_D166A) or 2 (PelA_h_, PelA_h_ E218A) independent experiments performed in duplicate with error bars indicating SE. **(b)** Pulmonary injury as measured by lactose dehydrogenase activity of the bronchoalveolar lavage fluid from BALB/c mice treated intratracheally or not with the indicated quantities of Sph3_h_ and sacrificed 7 days post treatment. Data represents the mean of at least *n* = 5, with error bars representing SE. **(c)** Fungal burden of neutropenic mice as determined by quantitative PCR following 4 days of infection with the indicated *A. fumigatus* strain with or without treatment with a single dose of 500 μg Sph3_h_. Data represents the mean of at least *n* = 12, from two independent experiments with error bars indicating SE. The * indicate a significant difference *p* < 0.01 for (a) and (c), and < 0.05 for (b), relative to untreated controls using a two-way ANOVA for (a) and a one-way ANOVA for (b) and (c) with a Dunnett’s multiple comparison test. **(d)** Histopathological analysis of lung sections of mice from (c) stained with Periodic acid Schiff reagent. Arrow indicates hyphal lesion. Scale bars = 20 μm.

### Intratracheal Sph3_h_ is well tolerated, and attenuates fungal virulence in an immunocompromised mouse model of pulmonary aspergillosis

Given the ability of Sph3_h_ to protect epithelial cells for over 24 hours, this hydrolase was selected for evaluation *in vivo.* BALB/c mice treated intratracheally with doses up to 500 μg of Sph3_h_ exhibited no signs of stress, weight loss or change in body temperature post-treatment **(Fig. S6a and b)**. Additionally, no significant increase in pulmonary injury or inflammation between treated and untreated mice were observed as measured by bronchoalveolar lavage lactate dehydrogenase activity and total pulmonary leukocyte populations **(Fig. 5b and Fig. S6c)**. Collectively these results suggest that a single intratracheal dose of Sph3_h_ is well tolerated by mice.

To determine the ability of Sph3_h_ to attenuate virulence of *A. fumigatus,* neutropenic BALB/c mice were infected intratracheally with *A. fumigatus* conidia with or without the co-administration of 500 μg of Sph3_h_. Four days after infection, mice infected with *A. fumigatus* and treated with Sph3_h_ had a significantly lower pulmonary fungal burden to untreated, infected mice as measured by both fungal DNA **(Fig. 5c)** and pulmonary galactomannan content **(Fig. S7)**. The fungal burden of the Sph3_h_-treated mice was similar to that observed with mice infected with the GAG-deficient hypovirulent strain *Δuge3* (20). Consistent with the fungal burden data, histopathologic examination of lung sections revealed the presence of fungal lesions in untreated, infected mice, but no detectable lesions in the lungs of infected mice treated with Sph3_h_, or those infected with conidia of the *Δuge3* mutant **(Fig. 5d)**. These findings suggest that Sph3_h_-mediated degradation of GAG can limit the growth of *A. fumigatus in vivo,* to the same degree as is observed with GAG-deficient organisms.

## Discussion

In this study, we demonstrate that the fungal glycoside hydrolase Sph3_h_ is able to degrade pre-formed *A. fumigatus* biofilms. This study is the first example of the use of a glycoside hydrolase to disrupt a fungal biofilm. Further, we establish that the glycoside hydrolase PelA_h_ displays activity against biofilms formed by organisms across different microbial kingdoms. Both glycoside hydrolases potentiated the penetration and activity of antifungal agents *in vitro,* exhibited no toxicity against mammalian cells and protected epithelial cells from *A. fumigatus-induced* damage. Pulmonary administration of Sph3_h_ was well tolerated and limited fungal growth in an immunocompromised mouse model, suggesting that these enzymes are promising therapeutic agents for the treatment of fungal disease.

The mechanism by which the biofilm matrix enhances *A. fumigatus* resistance to antifungals is poorly understood. The effect of hydrolase treatment on the antifungal sensitivity of *A. fumigatus* provides some insight into this question and establishes a role for GAG in biofilm-associated antifungal resistance. Multiple observations suggest that GAG enhances antifungal resistance by acting as a barrier to antifungal penetration of hyphae. First, glycoside hydrolase degradation of GAG enhanced the activity of multiple antifungals with different mechanisms of action. Second, the activity of posaconazole, but not voriconazole, was enhanced even though both azoles target the same enzyme, CYP51A. These hydrolases also display similar activity against azole-resistant and azole-sensitive strains. The cationic nature of GAG may explain the differential effects on voriconazole as compared with other antifungals. The GAG barrier would be predicted to be most effective against large, lipophilic or cationic antimicrobial agents, and thus therapeutic hydrolases may be most effective as adjuvants for lipophilic antifungals. Previous studies have reported that the enzymatic degradation of neutral *α*-glucans of *A. fumigatus* did not enhance susceptibility to antifungals (31), further supporting our hypothesis that exopolysaccharide charge plays a role in mediating antibiotic resistance. Similarly, hydrolysis of cationic Pel exopolysaccharide by PelA_h_ enhances the activity of the polycationic antibacterial colistin (22). Interestingly, degradation of biofilm-associated extracellular DNA (eDNA) has previously been reported to enhance *A. fumigatus* susceptibility to caspofungin and amphotericin B, though the effects on posaconazole and voriconazole susceptibility were not reported in the study (26). Recent work has suggested that Pel anchors eDNA within *P. aeruginosa* biofilms through charge-charge interactions (17). Given the similarities between Pel and GAG, it is possible that GAG-mediated binding of eDNA may also contribute to enhancing antifungal resistance.

The results of these studies add to an emerging body of evidence that fungal biofilms share structural (32–35) and functional (26, 36,37) similarity with those formed by pathogenic bacteria. The finding that glycoside hydrolases can display activity against the exopolysaccharides and biofilms of both fungi and bacteria provides the first evidence that these similarities could potentially be exploited for the development of therapeutics active against both organisms. Additionally, the similarity between the exopolysaccharides of *P. aeruginosa* and *A. fumigatus*, coupled with the interspecies activity of their glycoside hydrolases suggest the intriguing possibility that exopolysaccharide interactions may occur between organisms during multispecies biofilm formation. Co-colonization with *P. aeruginosa* and *A. fumigatus* is not uncommon in patients with chronic pulmonary disease such as cystic fibrosis (38). Although studies of the formation of mixed fungal-bacterial biofilms during pulmonary infection are limited, a recent study of patients with chronic lung disease reported that antibacterial therapy for *P. aeruginosa* was associated with a reduction in fungal colonization, suggesting the possibility of microbial cooperation (39). Further studies examining the role of cross-species exopolysaccharide and exopolysaccharide-modifying enzyme interactions are required to establish a role for cooperative biofilm interactions in pulmonary disease.

While PelA_h_ exhibited cross-species activity and disrupted pre-formed fungal biofilms, Sph3_h_ bound Pel, but was unable to disrupt established Pel-mediated biofilms. This difference in activity may reflect differences in the composition or conformation of each polysaccharide since GAG is a heteropolymer of GalNAc and galactose while Pel is comprised of GalNAc and GlcNAc. It is likely that these differences influence the ability of Sph3_h_ and PelA_h_ to hydrolyze the polymer. The inability of Sph3_h_ to degrade preformed *P. aeruginosa* biofilms may suggest that mature Pel adopts a configuration or undergoes post-synthetic modification that renders it incompatible with the catalytic active site of Sph3_h_ and resistant to cleavage. Detailed studies of these enzymes to determine the mechanisms underlying their differential activity against Pel will require purified polysaccharide, which is currently not available.

Both Sph3_h_ and PelA_h_ were found to be non-cytotoxic, and a single dose of intratracheal Sph3_h_ was well tolerated by BALB/c mice. Co-administration of Sph3_h_ with wild-type conidia to neutropenic mice greatly reduced fungal outgrowth within the lungs of these mice. Together these results provide proof-of-concept that the glycoside hydrolases can be used to improve the outcome of fungal infection, with minimal side effects and toxicity. These findings will pave the way for future work to evaluate the utility of these agents as antifungal therapeutics including detailed pharmacokinetic and toxicity studies, as well as the evaluation of these enzymes for the treatment of established fungal infections alone and in combination therapy with lipophilic antifungal agents such as posaconazole or amphotericin B.

## Methods

### Strains and culture conditions

Strains used in this study are detailed in Table S1 and detailed culture conditions are described in the **Supplementary Information (SI). Recombinant hydrolase expression and purification.** Hydrolases were expressed and purified as described previously (22, 23).

### Treatment of *A. fumigatus* with glycoside hydrolases

To visualize the effects of hydrolases on cell wall-associated GAG, hyphae were treated with recombinant hydrolases and stained with fluorescein-conjugated soybean agglutinin as previously described (23), with minor modifications. Hyphae were counterstained with a 1:1000 dilution of DRAQ5 (eBioscience) in phosphate buffered saline (PBS) for 5 min prior to paraformaldehyde (PFA) fixation. Complete image acquisition and processing methods can be found in the **SI**. To study the effects of hydrolases on biofilms, 10^4^ conidia were grown in Brian media in polystyrene, 96-well plates for 19 h at 37 °C and 5% CO_2_ and then treated with the indicated concentration of glycoside hydrolase in PBS for 1 h at room temperature. Biofilms were then gently washed, stained with 0.1% (w/v) crystal violet and de-stained with 100% ethanol for 10 min. The optical density of the de-stain fluid was measured at 600 nm (OD_600_).

### Scanning electron microscopy

Conidia were grown for 9 h in Dulbecco's Modified Eagle's Medium (DMEM) at 37 °C, 5% CO_2_ on glass coverslips, washed once with Ham's F- 12K (Kaighn's) Medium, and incubated with 500 in F-12K Medium with or without 0.5 μM hydrolase for 3 h at 37°C, 5% CO_2_. Coverslips were processed for scanning electron microscopy as previously described (20), and detailed in the **SI**.

### Treatment of *P. aeruginosa* with glycoside hydrolases

For biofilm disruption, static *P. aeruginosa* cultures were grown for 22 h at 30 °C, at which point the planktonic cells were aspirated and LB + 0.5% arabinose + 0.5 μM glycoside hydrolase was added for an additional 3 h. For the detection of Pel, samples were incubated with 30 μg/ml of fluorescein-conjugated *Wisteria fluoribunda* lectin for 2 h at 4 °C, fixed with 8% (w/v) PFA for 20 min at 4° C and imaged as detailed in the **SI**. The ability of hydrolases to disrupt established biofilms were studied as previously described (22).

### Culture supernatant production

*P. aeruginosa* cultures were grown at 30 °C for 24 h shaking at 200 rpm. Cultures were then centrifuged at 311 *x g* for 10 mins, and supernatants were filtered using 0.44 μm syringe filters. Culture supernatants were stored at −20 °C until use.

### Hydrolase binding quantification

Undiluted culture supernatants were incubated on Immunolon^®^ 2HB high-binding 96 well microtiter plates overnight at 4 °C. Wells were washed 3X with Wash Buffer (PBS + 0.05% (v/v) Tween-20) and blocked for 30 min at 4 °C in Blocking Buffer (1% (w/v) Bovine Serum Albumin in Wash Buffer). Wells were washed 1X with Wash Buffer and incubated with the indicated concentrations of Sph3_h_ diluted in Blocking Buffer for 3 h at 4 °C. Wells were washed 3X with Wash Buffer and incubated with polyclonal rabbit anti-Sph3_h_ (custom-produced by Cedarlane Laboratories) diluted 1/100 in Blocking Buffer for 1.5 h at 4 °C. Wells were then washed 3X with Wash Buffer and incubated with donkey-anti-rabbit secondary antibody conjugated to horseradish peroxidase diluted 1/2000 in Blocking Buffer for 1 h at 4 °C. Wells were then washed 4X with Wash Buffer and incubated with TMB substrate (ThermoFisher®) for 15 mins at room temperature. The reaction was stopped with the addition of 2 N H_2_SO_4_ and absorbance read at 450 nm with a 570 nm correction.

### Effects of glycoside hydrolases on antifungal susceptibility of *A. fumigatus.*

Fungal biofilms were prepared in tissue culture treated 24 well plates in RPMI 1640 medium (Life Technology) buffered with MOPS (3-(N-Morpholino) Propane-Sulfonic Acid) (Fisher) (RPMI-MOPS) for 9 h at 37 °C, 5% CO_2_. Serial dilutions of antifungal compounds with or without 0.5 μM of Sph3_h_ or PelA_h_ were added to wells and the plates incubated at 37 °C and 5% CO_2_ for 15 h. Fungal viability was measured using the sodium 3'-[1-[(phenylamino)-carbony]-3,4-tetrazolium]-bis(4-methoxy-6-nitro)benzene-sulfonic acid hydrate (XTT) metabolic assay as described previously (40). The concentration of antifungal resulting in a 50% decrease in viability (MIC_50_) was used as a measure of antifungal effect.

### Fluorometric quantification of hyphal uptake of BDP-PCZ

2.5x10^4^ conidia of red fluorescent protein (RFP)-expressing *A. fumigatus* were grown in a 96-well black, clear-bottom plate for 8 h at 37 °C, 5% CO_2_. Hyphae were treated with 1 μM of Sph3_h_ in PBS for 90 min at 37 °C, 5% CO_2_ then exposed to 2 μg/mL BDP-PCZ for 10 mins. The plate was then read using an Infinite M1000 fluorescent plate reader with excitation wavelengths of 532 and 488 nm for RFP and BDP-PCZ, respectively. Background fluorescence was subtracted from both RFP and BDP-PCZ signals and the BDP-PCZ signal was then normalized to total RFP fluorescence for each well.

### Effects of glycoside hydrolases on *A. fumigatus-induced* epithelial cell damage

A549 pulmonary epithelial cell damage by *A. fumigatus* was tested using the ^51^Cr release assay as previously described (20, 30). Recombinant hydrolases were added to the A549 cultures at the time of infection at a final concentration of 0.5 μM.

### Characterization of pulmonary damage by Sph3_h_

All procedures involving mice were approved by the Animal Care Committees of the McGill University Health Centre. Female BALB/c mice 5-6 weeks of age were anaesthetized with isoflurane and administered a single endotracheal injection of 500 μg Sph3_h_ in 50 μL PBS and monitored daily for 7 days. Mice were then euthanized by CO_2_ overdose and their airways lavaged with 1 mL PBS that was administered and collected through a needle inserted in the trachea. A total of 2 lavages were performed and pooled. The presence of LDH in the BAL fluid was used as an indicator of pulmonary damage; LDH activity was measured in the fluid using a commercial assay (Promega), as per manufacturer’s instructions.

### Effects of Sph3_h_ in a severely immunocompromised mouse model of invasive pulmonary aspergillosis

Mice were immunosuppressed with cortisone acetate and cyclophosphamide as previously described (20, 41). Mice were infected with an endotracheal injection of 5x10^3^ *A. fumigatus* conidia, resuspended in either PBS alone, or in combination with 500 μg of Sph3_h_. Mice were monitored daily and moribund animal were euthanized. At 4 days post infection mice were euthanized and their lungs were harvested. For fungal burden analysis, lungs were homogenized in 5 mL PBS containing protease inhibitor cocktail (Roche), and aliquots were stored at −80°C until use. Pulmonary fungal burden was determined as previously described (15), and detailed in the **SI**. For histological examination, lungs were inflated with 10% buffered formalin (Fisher Scientific) and immersed in formalin overnight to fix. Lungs were then embedded in paraffin and 4 μm thick sections were stained with periodic acid Schiff (PAS) stain.

## Acknowledgements

Research described in this paper is supported by operating grants from the *Canadian Institutes of Health Research* (CIHR) (#81361 to P.L.H. and D.C.S; #123306 to D.C.S.; and #43998 to P.L.H) and Cystic Fibrosis Canada (CFC) (to D.C.S. and P.L.H). B.D.S has been supported by graduate scholarships from CFC and CIHR. N.C.B has been supported in part by graduate scholarships from the *Natural Sciences and Engineering Research Council of Canada,* Mary H. Beatty, and Dr. James A. and Connie P. Dickson Scholarships from the University of Toronto, CFC, and The Hospital for Sick Children. P.B. has been supported in part by a CFC postdoctoral fellowship and a Banting Fellowship from CIHR. P.L.H. is the recipient of a Canada Research Chair. D.C.S is supported by a Chercheur-Boursier Award from the *Fonds de Recherche Quebec Santé.*

## References

1. Bajwa SJ, Kulshrestha A (2013) Fungal infections in intensive care unit: Challenges in diagnosis and management. Ann Med Health Sci Res 3(2):238.

2. Peleg AY, Hooper DC (2010) Hospital-acquired infections due to gram-negative bacteria. N Engl J Med 362(19):1804–1813.

3. Vincent JL, et al. (1995) The prevalence of nosocomial infection in intensive care units in Europe. Results of the European Prevalence of Infection in Intensive Care (EPIC) Study. EPIC International Advisory Committee. JAMA 274(8):639–644.

4. Maltezou HC (2009) Metallo-β-lactamases in Gram-negative bacteria: introducing the era of pan-resistance? International Journal of Antimicrobial Agents 439 33(5):405.e1–405.e7.

5. Brown GD, et al. (2012) Hidden killers: human fungal infections. Science Translational Medicine 4(165):165rv13.

6. Verweij PE, et al. (2015) International expert opinion on the management of infection caused by azole-resistant Aspergillus fumigatus. Drug Resist Updat 21-22:30–40.

7. Mann EE, Wozniak DJ (2012) Pseudomonas biofilm matrix composition and niche biology. FEMS Microbiology Reviews 36(4):893–916.

8. Gravelat FN, et al. (2010) Aspergillus fumigatus MedA governs adherence, host cell interactions and virulence. Cellular Microbiology 12(4):473–488.

9. Sheppard DC (2011) Molecular mechanism of Aspergillus fumigatus adherence to host constituents. Current Opinion in Microbiology 14(4):375–379.

10. Høiby N, Bjarnsholt T, Givskov M, Molin S, Ciofu O (2010) Antibiotic resistance of bacterial biofilms. International Journal of Antimicrobial Agents 35(4):322–332.

11. Seidler MJ, Salvenmoser S, Muller FMC (2008) Aspergillus fumigatus Forms Biofilms with Reduced Antifungal Drug Susceptibility on Bronchial Epithelial Cells. Antimicrobial Agents and Chemotherapy 52(11):4130–4136.

12. Alhede M, Bjarnsholt T, Givskov M, Alhede M (2014) Pseudomonas aeruginosa biofilms: mechanisms of immune evasion. Adv Appl Microbiol 86:1–40.

13. Lee MJ, et al. (2015) The Fungal Exopolysaccharide Galactosaminogalactan Mediates Virulence by Enhancing Resistance to Neutrophil Extracellular Traps. PLoS Pathog 11(10):e1005187.

14. Fontaine T, et al. (2011) Galactosaminogalactan, a new immunosuppressive polysaccharide of Aspergillus fumigatus. PLoS Pathog 7(11):e1002372.

15. Lee MJ, et al. (2016) Deacetylation of Fungal Exopolysaccharide Mediates Adhesion and Biofilm Formation. mBio 7(2):e00252–16.

16. Franklin MJ, Nivens DE, Weadge JT, Howell PL (2011) Biosynthesis of the Pseudomonas aeruginosa Extracellular Polysaccharides, Alginate, Pel, and Psl. Front Microbiol 2:167.

17. Jennings LK, et al. (2015) Pel is a cationic exopolysaccharide that cross-links extracellular DNA in the Pseudomonas aeruginosa biofilm matrix. Proc Natl Acad Sci USA 112(36):11353–11358.

18. Colvin KM, et al. (2011) The Pel and Psl polysaccharides provide Pseudomonas aeruginosa structural redundancy within the biofilm matrix. Environmental Microbiology 14(8):1913–1928.

19. Pritt B, O'Brien L, Winn W (2007) Mucoid Pseudomonas in Cystic Fibrosis. American Journal of Clinical Pathology 128(1):32–34.

20. Gravelat FN, et al. (2013) Aspergillus galactosaminogalactan mediates adherence to host constituents and conceals hyphal β-glucan from the immune system. PLoS Pathog 9(8):e1003575.

21. Yang L, et al. (2012) Polysaccharides serve as scaffold of biofilms formed by mucoid Pseudomonas aeruginosa. FEMS Immunology & Medical Microbiology 65(2):366–376.

22. Baker P, et al. (2016) Exopolysaccharide biosynthetic glycoside hydrolases can be utilized to disrupt and prevent Pseudomonas aeruginosa biofilms. Science Advances 2(5):e1501632.

23. Bamford NC, et al. (2015) Sph3 is a glycoside hydrolase required for the biosynthesis of galactosaminogalactan in Aspergillus fumigatus. Journal of Biological Chemistry 290(46):27438–27450.

24. Khan W, et al. (2010) Aminoglycoside resistance of Pseudomonas aeruginosa biofilms modulated by extracellular polysaccharide. Int Microbiol 13(4):207–212.

25. Colvin KM, et al. (2011) The Pel Polysaccharide Can Serve a Structural and Protective Role in the Biofilm Matrix of Pseudomonas aeruginosa. PLoS Pathog 7(1):e1001264.

26. Rajendran R, et al. (2013) Extracellular DNA release acts as an antifungal resistance mechanism in mature Aspergillus fumigatus biofilms. Eukaryotic Cell 12(3):420–429.

27. Rajendran R, et al. (2011) Azole Resistance of Aspergillus fumigatus Biofilms Is Partly Associated with Efflux Pump Activity. Antimicrobial Agents and Chemotherapy 55(5):2092–2097.

28. Mowat E, et al. (2008) Phase-dependent antifungal activity against Aspergillus fumigatus developing multicellular filamentous biofilms. Journal of Antimicrobial Chemotherapy 62(6):1281–1284.

29. Campoli P, et al. (2013) Pharmacokinetics of posaconazole within epithelial cells and fungi: insights into potential mechanisms of action during treatment and prophylaxis. Journal of Infectious Diseases 208(10):1717–1728.

30. Bezerra LML (2004) Interactions of Aspergillus fumigatus with endothelial cells: internalization, injury, and stimulation of tissue factor activity. Blood 103(6):2143–2149.

31. Beauvais A, et al. (2007) An extracellular matrix glues together the aerial-grown hyphae of Aspergillus fumigatus. Cellular Microbiology 9(6):1588–1600.

32. Martins M, et al. (2009) Presence of Extracellular DNA in the Candida albicans Biofilm Matrix and its Contribution to Biofilms. Mycopathologia 169(5):323–331.

33. Krappmann S, Ramage G (2013) A sticky situation: extracellular DNA shapes Aspergillus fumigatus biofilms. Front Microbiol 4:159.

34. Fuxman Bass JI, et al. (2010) Extracellular DNA: A Major Proinflammatory Component of Pseudomonas aeruginosa Biofilms. The Journal of Immunology 184(11):6386–6395.

35. Vilain S, Pretorius JM, Theron J, Brozel VS (2009) DNA as an Adhesin: Bacillus cereus Requires Extracellular DNA To Form Biofilms. Applied and Environmental Microbiology 75(9):2861–2868.

36. Martins M, Henriques M, Lopez-Ribot JL, Oliveira R (2011) Addition of DNase improves the in vitro activity of antifungal drugs against Candida albicans biofilms. Mycoses 55(1):80–85.

37. Chiang WC, et al. (2013) Extracellular DNA Shields against Aminoglycosides in Pseudomonas aeruginosa Biofilms. Antimicrobial Agents and Chemotherapy 57(5):2352–2361.

38. Paugam A, et al. (2010) Characteristics and consequences of airway colonization by filamentous fungi in 201 adult patients with cystic fibrosis in France. Med Mycol 48 Suppl 1(O1):S32–S36.

39. Baxter CG, et al. (2013) Intravenous antibiotics reduce the presence of Aspergillus in adult cystic fibrosis sputum. Thorax 68(7):652–657.

40. Pierce CG, et al. (2008) A simple and reproducible 96-well plate-based method for the formation of fungal biofilms and its application to antifungal susceptibility testing. Nat Protoc 3(9):1494–1500.

41. Sheppard DC, et al. (2004) Novel inhalational murine model of invasive pulmonary aspergillosis. Antimicrobial Agents and Chemotherapy 48(5):1908–1911.

42. Brannon MK, et al. (2009) Pseudomonas aeruginosa Type III secretion system interacts with phagocytes to modulate systemic infection of zebrafish embryos. Cellular Microbiology 11(5):755–768.

